## Supplemental Information for "Fierce selection and interference in B-cell repertoire response to chronic HIV-1"

### Contents


**1 BCR repertoire data** **1**

**2 Inference of lineage phylogenies** **2**

**3 Inference of selection from lineage tree statistics** **3**

**4 Selection and clonal interference likelihood ratios** **5**

**5 Robustness of selection statistics** **10**

**6 Simulations** **10**


### 1 BCR repertoire data

**HIV patients.** We analyze B-cell repertoire data from 6 HIV patients from ref. [1] with raw sequence reads accessible from the European Nucleotide Archive under study accession numbers, ERP009671 and ERP000572. We study the repertoire data in two untreated HIV patients with run accession numbers, ERR769563 - 9569 (patient 1) and ERR162588 - 2595 (patient 2) and in four patients with ART interruption at week 48, ERR769570 - 9576 (patient 3), ERR769541 - 9547 (patient 4), ERR769548 - 9554 (patient 5), ERR769534 - 9540 (patient 6). The data covers $\sim 2.5$ years of study with 6-8 sampled time points per patient; see Table S1 for details.

The B-cell repertoire sequences consist of 150bp non-overlapping paired-end reads (Illumina MiSeq), with one read covering much of the V gene and the other read covering the area around the CDR3 region and the J gene. The average gap length between the two reads is approximately 35 nucleotides. The gapped region overlaps partly with FWR3 of BCRs. Further details on the experimental protocol are discussed in ref. [1]. For the initial processing of the raw reads we use pRESTO [2] (version 0.5.2) with the following steps: we filter sequences for quality ($> 32$) and length ($> 100$). The small fraction of the paired end reads that overlap are assumed to be anomalous, and are discarded from the analysis. We assemble the paired reads by aligning against the IMGT reference database of V genes [3], such that an appropriate size gap is inserted between the non-overlapping paired reads. Duplicate sequences are collapsed into unique sequences.

The sequences contained a large number of singletons, that is sequences with no duplicates. We use a statistical approach to establish whether a singleton is a "true" independent BCR or an erroneous duplicate of another BCR, which has already been accounted for. We argue that sequences with few differences from non-singletons are more likely to have appeared due to sequencing errors. With an R script, we calculate the minimum Hamming distance of each singleton to any non-singleton, $H_0$ (Fig. S1a). The distribution of $H_0$ in the data $W(H_0)$ is a mixture of a Poisson distribution $\text{Poiss}(H_0; \nu)$ with parameter $\nu$ , related to sequencing errors at small values of $H_0$, and a Gamma distribution $\text{Gam}(H_0; \alpha, \beta)$ with parameters $\alpha$, $\beta$, describing the remaining $H_0$'s that reflect the distances among the sampled BCRs. The mixture distribution can be written as $W(H_0) = (1-\lambda)\,\text{Poiss.}(H_0; \nu) + \lambda\,\text{Gam.}(H_0; \alpha, \beta)$, with a hidden variable $\lambda$. We infer maximum likelihood estimates for the parameters of the distributions, $\nu$, $\alpha$, $\beta$ and their relative weight $\lambda$. The probability for a singleton with a Hamming distance $H_0$ to be a true independent BCR sequence and not to be generated by sequencing error follows, $\rho(H_0) = \lambda\,\text{Gam.}(H_0; \alpha, \beta)/W(H_0)$. The probability density $\rho(H_0)$ is a sigmoid function in all patients, indicating that sequences with few differences to the existing non-singletons are more likely to have appeared due to sequencing errors (insert in Fig. S1a). We use a conservative estimate of $\rho(H_0)$ and discard all singletons with $H_0 < 5$ (i.e., those with $\rho(H_0) < 1$ ) from our analysis. Due to lack of barcoding for individual molecules, we only use the unique BCR sequences for analysis and do not incorporate the information on the multiplicity of each sequence.

**Healthy individuals.** We analyze memory B-cell repertoire data of 3 individuals published in ref. [4]: https://clients.adaptivebiotech.com/pub/robins-bcell-2016. The published data in healthy individuals is already pre-processed for quality control and corrected for sequencing error. In particular, the singletons due to sequencing errors have been already discarded from the dataset. Fig. S1b shows the distribution for the minimum Hamming distance of each singleton to any non-singleton, $W(H_0)$, in the healthy repertoire data. Maximum likelihood inference of a mixture model for $W(H_0)$, as described above for repertoires in HIV patients, suggests that BCRs in healthy repertoire data are not likely to have been generated by sequencing errors (insert in Fig. S1b).

**BCR annotation.** In both datasets, we annotate the BCR repertoire sequences of each individual (pooled time points) by Partis (version 0.11.0) [5]. Partis uses very large amounts of memory, so the initial (cache-parameters) stage is run on a subset of 200,000 random sequences, and the annotation stage is run on the full set of sequences. We process the output of Partis in R, which includes the estimated V gene/allele, J gene/allele, location of the CDR3 region, and an inferred naive sequence (germline before hyper-mutation). Sequences which have indels outside of the CDR3 are discarded. We partition the sequences into two groups: productive BCRs, which are in-frame and have no stop codons, and the unproductive BCRs. The sequences are further annotated by processing the inferred naive sequences with MiXCR [6, 7], which gives the CDR1, CDR2 and framework regions. The sequence reads in healthy individuals [4] are shorter than in HIV patients [1] and do not extend to CDR1/2.

**Lineage reconstruction.** To identify BCR lineages, we first group sequences by the assigned V gene, J gene and CDR3 length, and then used single linkage clustering with a threshold of 90% Hamming distance. A similar threshold has been previously suggested by ref. [8] to identify BCR lineages. Clusters of small size ($< 20$) are discarded from our analysis. For each cluster, there may be multiple inferred naive sequences, as this is an uncertain estimate, and the most common naive sequence is chosen to be the outgroup for genealogy reconstruction. See Table S1 for detailed statistics of BCR lineages in each individual.

**Unproductive BCRs.** Due to a larger sequencing depth in healthy individuals, we are able to reconstruct relatively large unproductive BCR lineages. Unproductive sequences are BCRs that are generated, but due to a frameshift or insertion of stop codons, are never expressed. These BCRs reside with productive (functional) BCRs in a nucleus and undergo hypermutation during B-cell replication, and therefore, provide a suitable null expectation for somatic evolution during affinity maturation.

#### 2 Inference of lineage phylogenies

**Lineage genealogy reconstruction.** For each lineage and its aligned sequences we reconstruct its underlying genealogical tree. We use FastTree [9] to construct the initial tree by maximum parsimony. We use this tree as seed for the maximum likelihood construction of the phylogeny with RAxML [10], using the GTRCAT substitution model. In the last step of tree topology reconstruction, we use the GTRGAMMA substitution model to optimize sequence divergence along the tree (i.e., branch lengths). We use a maximum likelihood approach to reconstruct nucleotide sequences of internal nodes on the tree [11]. We do not include the positions with gaps in the multi-sequence alignment in inference of tree topology and the nucleotide mutations along the tree.

We use the inferred naive sequence (germline) as the outgroup of the genealogy. The root of the tree may be some mutations away from the last common ancestor of the sampled sequences. This may be due to a number of initial rounds of hypermutation prior to secretion of the first selected B-cell, or alternatively, due to incorrect assignment of the germline allele during annotation; a fraction of V, D and J alleles circulating in the human population are missing from the existing reference datasets like IMGT [3]. In order to minimize the effect of such allele mis-assignments, we discard the mutations that separate the inferred germline sequence and the last common ancestor (root) of the tree from our analysis (i.e., mutations common to all sequences).

**Inference of branching time along a phylogeny.** To characterize the branch length statistics of lineages in units of coalescence time (used in Fig. 2 and Fig. S2b), we use a maximum-likelihood approach and a probabilistic model to annotate internal nodes of a tree with times of occurrence, given the topology of the tree and the mutations on the branches. Internal nodes represent replication events, which may carry new mutations assigned to branches by the ancestral sequence reconstruction procedure. The model can also correct observation times of external nodes on the tree, given sufficient evidence. The model assumes a tree with $n$ nodes, $[o_1, \ldots, o_m, i_{m+1}, i_n]$, where $o_k$ are the sampled sequences in the leaves of the tree, and $i_k$ are the internal nodes of the tree. The observed nodes are annotated with their sampling times, $\mathbf{T} = [T_1, \ldots, T_m]$. Branches of the tree are annotated with their mutational distances, $\mathbf{d} = [d_1, \ldots, d_n]$. We only consider synonymous mutations for computation of mutational distances. The model estimates mutation rate, $\mu$ and the times $\mathbf{t} = [t_1, \ldots, t_n]$ of the nodes, by maximizing the likelihood:

$$P(\mathbf{d}, \mathbf{T}|\mathbf{t}, \mu) = \prod_{k=1}^{m} \frac{1}{\sigma\sqrt{2\pi}} \exp\left[-\frac{(t_k - T_k)^2}{2\epsilon}\right] \prod_{j=1}^{n} \frac{(\mu\tau_j)^{d_j}}{d_j!} \exp[-\mu\tau_j], \quad (1)$$

where $\tau_j = t_j - t_{\mathcal{A}(j)}$, the time difference between a node in the tree and its parent, is constrained to be positive. The model assumes Gaussian measurement error with standard deviation $\sigma$ of the sampling time for the observed nodes, and the Poisson model for mutations on tree branches, with rate $\mu$. To limit the search space of the optimization algorithm we constrain the times $\mathbf{t}$ to be discrete; the units of time used in our data are weeks. Here we set $\sigma = 5$. The optimization is solved by an iterative procedure in which times of nodes are changed by one unit in the direction of score increase, until convergence.

#### 3 Inference of selection from lineage tree statistics

We compare genealogies of B-cell lineages in HIV patients with those of healthy individuals to characterize the evolutionary selection during affinity maturation in response to chronic infection. The structure of genealogies has been linked to evolutionary modes in a population [12]. Rapid evolution under positive selection leads to skewed tree topologies and elongated terminal branches [11, 13–15], compared to neutral evolution [12, 16] (Fig. S2a). Selection in BCR repertoires has recently been quantified in response to influenza vaccination, using BCRs sampled over a relatively short time span post-vaccination with a peak response at day 7 [17]. Compared to the Horns et al study [17], we study populations on longer time-scales that are better described as well-mixed populations. For these reasons our results are less sensitive to the effects of spatial segregation and additional biases resulting from B-cells exiting germinal centers.

**Asymmetric tree branching.** We characterize the asymmetry of trees by the branching imbalance of the last common ancestor at the root of the tree - the last common ancestor may be a number of mutations away from the germline progenitor. We define the weight of each node in a tree by the number of leaves (terminal nodes) within its clade; see Fig. S2a [15]. The weight of the last common ancestor $w_{\text{anc..}}$, i.e., the total number of leaves in a tree, and its daughters, $w_{D_1}$, $w_{D_2}$, are indicated in the simulated trees of Fig. S2a. In rapid evolution under selection, the first branching event produces highly imbalanced sub-clades, and hence, extreme values for the weight of the first daughter nodes [15]. In contrast, neutral evolution predicts a uniform distribution of tree weights of ancestral sub-clades. Fig. 2a shows a U-shaped distribution of relative weights of daughters to the ancestor $w_D/w_{\text{anc.}}$ in lineages reconstructed from BCRs in HIV patients and in healthy individuals, indicating intra-lineage selection during affinity maturation.

**Terminal branch statistics.** In lineages under selection, the descendent sequences (leaves of a tree) are likely to coalesce to an ancestor with high fitness, resulting in long terminal branches in a tree, in contrast to the exponentially distributed branch lengths in neutrality (Fig. S2a) [14, 15] — branch length statistics are estimated in units of coalescence time, rather than sequence Hamming distance; see the paragraph on inference of branching times along a phylogeny. In Fig. 2b we compare the ratio of mean terminal branch length of a lineage to the averaged length of all the branches in a lineage. The distribution of this branch length ratio in HIV patients show an excess of lineages with relatively long terminal branches, compared to the expected distribution for simulated neutral lineages of the same size (Kingman's coalescence); see the paragraph on simulated trees. A similar trend with a weaker signal is seen in healthy individuals (Fig. 2b). To make sure that the strong signal in HIV patients is not only due to sampling of the repertoire over multiple time points, we repeat the same analysis on the subset of lineages which are sampled in only one time point. Fig. S2b shows a comparable over-representation of long terminal branches in this subset.

The branch length ratio (Fig. 2 and Fig. S2) is a dimensionless quantity, and hence, is comparable across datasets with different sampling times. The consistency between the lineage trees sampled in a single and multiple time points (Fig. S2) indicates that the inferred time statistics is robust for such analysis. Nonetheless, this inference provides only an effective time approximation for the rise of mutations, which we do not rely upon in the other selection inference procedures, besides the branch length statistics.

**Site frequency spectrum (SFS).** The SFS is the probability density $f(\nu)$ of observing a derived mutation (allele) with a given frequency $\nu$ in a lineage. A mutation that occurs along the phylogeny of a lineage forms a clade and is present in all the descendent nodes (leaves) of its clade (see Fig. S2a). The SFS carries information about the shape of the phylogeny, including both the topology and the branch lengths. In neutrality, mutations rarely reach high frequency, and hence, the SFS decays monotonically with allele frequency as, $f(\nu) \sim \nu^{-1}$ [16]. In phylogenies with skewed branching, many mutations reside on the larger sub-clade following a branching event, and hence, are present in the majority of the descendent leaves on the tree. The SFS of such lineages is often non-monotonic with an upturn in the high frequency part of the spectrum and a steeper drop ($\sim \nu^{-\beta}$ with $\beta > 1$) in the low frequency part of the spectrum [14].

To identify the targets of selection, we classify mutations based on the region they occur in. BCRs are made up of the three immunologically important complementarity-determining regions (CDRs) [18], CDR1, CDR2 and CDR3 and the remaining part of the V and J genes referred to as the framework region (FWR). We evaluate the frequency of a given mutation on a tree as the fraction of the observed sequences (leaves) which carry that mutation, which is approximately equal to the size of the its descendent clade relative to the whole tree (up to the effect of back mutations). Estimating the SFS based on the mutations inferred on a tree rather than just the variations among the observed sequences allows us to more accurately identify the derived alleles. In addition, the structure of the tree has information about the sequences which are not sampled, sequenced or clustered within a lineage.

Fig. 2c shows the SFS pooled across lineages in HIV patients and in healthy individuals; mutation frequencies are estimated within each lineage. We see a signifiant upturn of the SFS polarised on non-synonymous mutations in pathogen-engaging CDR3 regions; this signal of selection is strongest in HIV patients with an order of magnitude increase in the high end of the spectrum (Fig. 2c). The SFS polarised on mutations in other regions show steeper drop in the low frequency side of the spectrum compared to neutral expectation. Fig. S3 shows the SFS of the unproductive BCR lineages in healthy individuals, with a comparable steep drop in low frequencies. A similar pattern in the SFS has recently been reported for lineages of B-cell repertoires following influenza vaccination [17].

In order to estimate the significance of the upturn in the shape of the SFS, we simulated 100 realizations (numerical experiments) of lineages under neutral evolution (i.e., Kingman's coalescent) for each dataset (i.e., HIV patients, productive and unproductive lineages in healthy individuals); see the paragraph on Simulations. Each realization contains trees of the same sizes as in the corresponding experimental dataset. We estimate the SFS for each realization separately and show the span and confidence level for the SFS as a function of the mutation frequencies (grey area in Fig. 2, Fig. S3 ). Expectantly, the estimate of the SFS is noisier at the high-frequency end of the spectrum, where fewer mutations are present, and is noisiest for unproductive BCRs with only 131 lineages. To quantify the significance of the high-frequency upturn in the SFS, we compare the test statistics, $\delta f = f(\nu \to 1) \Big/ \min[f(\nu)]$ in real data, polarized on different types of mutations, to the simulated realizations of the SFS. The upturn is significant for non-synonymous CDR3 mutations and for synonymous mutations in productive lineages in HIV patients and in healthy individuals with p-value $< 0.03$[1]. In HIV patients, the upturn of the SFS polarized on the amino acid changes in CDR1/2 and FWR are insignificant, respectively with p-values: $0.26$ and $0.48$. In healthy individuals, the upturn is insignificant for FWR changes in productive lineages (p-value= $0.26$) and for all mutation types in unproductive lineages (with p-values in CDR3: $0.13$, FWR: $0.53$, synonymous: $0.45$).

While the upturn of the SFS is often used as a standard signal for selection in population genetics, one should be very cautious with the interpretation of the SFS signal. The tests based on the allele frequencies have in general low power in distinguishing between hitchhiking under selection or out-of-equilibrium effects due to population structure in neutrality. For example, demographic models could produce a similar pattern of the SFS with a high-frequency upturn [19, 20] which could be interpreted as a selective sweep using Fay and Wu's H-statistic [21]. This could in particular be misleading when applied to expanding B-cell populations in response to an acute infection or vaccination.

For analysis of lineage tree statistics, i.e., the weight imbalance and terminal branch statistics, we only rely on the relatively large lineages with size ($> 50$) leaves. To avoid the noisy mutation frequency estimates, we estimated SFS from trees with size $> 100$ (see Table S1 for detailed statistics).

#### 4 Selection and clonal interference likelihood ratios

**Selection likelihood ratio.** Hypermutations during affinity maturation create new clades within a lineage. The frequency $x$ of these clades change over time, as shown by the schematic in Fig. 3a. A mutation under positive (or negative) selection should reach a higher (lower) frequency than a neutral mutation. Many population genetics tests, such as the McDonnell-Kreitman test for positive selection [22], rely on a comparison between statistics of substitutions (i.e., mutations that fix within a population) and the circulating polymorphisms within species. Unlike phylogenies based on species divergence, B-cell lineages form genealogies with many mutations that rise to intermediate frequencies as polymorphisms but often do not fix within a lineage. Here, instead of relying on the substitution statistics, we use the history of polymorphisms to quantify selection in B-cell lineages. In particular, we estimate the frequency propagator $G(x)$ [23] as the likelihood that a new mutation (allele) appearing in a lineage reaches frequency $x$ at some later time within a lineage (see schematic in Fig. 3a).

To estimate intra-lineage selection, we compare the likelihood of an amino acid changing non-synonymous mutation reaching a given frequency $x$ at any point in its time trace, $G(x) \equiv n(x)/N$, to that likelihood for a synonymous mutation in the same lineage, $G_0(x) \equiv n_0(x)/N_0$. Here, $n(x)$ and $n_0(x)$, respectively, denote the number of non-synonymous and synonymous mutations that reach frequency $x$ (at any time in general) and $N$ and $N_0$ denote the total number of such mutations (Fig. 3a) [23]. We determine the selection selection likelihood ratio as,

[1] test statistic was more extreme than all 100 bootstraps

$$g(x) = \frac{G(x)}{G_0(x)}. \tag{2}$$

Due to heterogeneity and context dependence of mutation rates in different regions of BCRs, we evaluate the likelihood ratio separately for each region, namely the CDR3, CDR1 & CDR2 (pooled together) and framework regions (FWR). In the Fig. S10 we show the robustness of the region-specific selection likelihood ratio with respect to such mutational biases.

**Interference likelihood ratio (time-ordered selection).** Clonal interference among beneficial mutations on different genetic backgrounds is a characteristic of evolution in asexual populations. In the absence of clonal interference, beneficial mutations can readily fix in a population after they rise to intermediate frequencies, beyond which stochastic effects cannot impact their fate [24]. Clonal interference reduces the efficacy of selection, resulting in a quasi-neutral regime of evolution [25].

To examine the amount of clonal interference among BCRs within a lineage, we consider time ordered selection propagators (interference propagators) indicating the likelihood that a mutation reaches frequency $x$ and later goes extinct, $H(x) = G(x) \times G(0|x)$ [23]. Here, $G(0|x)$ is the conditional probability that a mutation trajectory decays to frequency 0 given that it starts from frequency $x$; see the schematic in Fig. 3a. We estimate the interference likelihood ratio by comparing the probability of a non-synonymous mutation to reach a frequency $x$ and later go extinct $H(x)$ to the same scenario for synonymous mutations, $H_0(x)$ (Fig. 3a),

$$h(x) = \frac{H(x)}{H_0(x)} \equiv \frac{G(x) \times G(0|x)}{G_0(x) \times G_0(0|x)}. \tag{3}$$

**Quantifying selection and interference from likelihood ratios.** Assuming that mutations occur independently in a lineage (i.e., an independent site model), we can analytically characterize the functional dependence of the likelihood ratios on the selection coefficients of the sites. The probability $G(x|x_i, t)$ that an allele with a starting frequency $x_i$ reaches a frequency $x$ by time $t$ satisfies the backward Kimura's equation [26],

$$\frac{\partial}{\partial t} G(x|x_i, t) = \left[\frac{1}{2N} x(1-x)\frac{\partial^2}{\partial x^2} + \sigma(t) x(1-x)\frac{\partial}{\partial x}\right] G(x|x_i, t), \tag{4}$$

with the boundary conditions $G(x|0) = 0$ and $G(x_i|x_i) = 1$. Here, we assume that $\sigma(t)$ is the selection coefficient with a fluctuating strength around a mean value, $\sigma(t) = \overline{\sigma} + \eta(t)$, where $\eta$ is an uncorrelated Gaussian random variable with mean 0 and variance $\langle \eta(t)\eta(t')\rangle = \nu\delta(t-t')$. We interpret these fluctuations as *micro-evolutionary* changes in the preference and relative fitness (i.e., selection) of the focal allele due to e.g., spontaneous changes in the state of the other linked loci. This provides an effective model for the short-time dynamics of an allele within a lineage. However, it does not fully capture all dynamical features of evolution with clonal interference, including *macro-evolutionary* (i.e., long term) correlations in environmental fluctuations due to the slow dynamics and rise of the other alleles.

The stationary state solution of the probability for an allele to reach frequency $x$ during its lifetime follows,

$$0 = \left[\left(x(1-x) + vx^2(1-x)^2\right)\frac{\partial^2}{\partial x^2} + \left(\overline{s}x(1-x) + vx(1-x)(1-2x)\right)\frac{\partial}{\partial x}\right] G(x|x_i), \tag{5}$$

where $s = 2N\sigma$ and $v = 2N\nu$ are rescaled selection coefficient and fluctuation strengths, respectively. The solution to the stationary state selection propagator follows [27],

$$G(x|x_i;\bar{s},v) = \frac{1-\left|\frac{1-x_i/a_+}{1-x_i/a_-}\right|^{\lambda(\bar{s},v)}}{1-\left|\frac{1-x/a_+}{1-x/a_-}\right|^{\lambda(\bar{s},v)}}, \tag{6}$$

with $a_\pm = \frac{1}{2}[1 \pm \sqrt{1+4/v}]$ and $\lambda(\bar{s},v) = \bar{s}/(v\sqrt{1+4/v})$.

In the absence of fluctuations ($v \to 0$ and $\bar{s} \equiv s$), the selection likelihood approaches the known limit [28],

$$G(x|x_i;s,v=0) = \frac{1-e^{-sx_i}}{1-e^{-sx}} \tag{7}$$

In the limit of small initial (entry) frequencies $x_i \ll 1$, the conditional probability can be approximated by,

$$G(x|\overline{x_i};\bar{s},v) = \frac{\bar{s}\,\overline{x_i}}{1-\left|\frac{1-x/a_+}{1-x/a_-}\right|^{\lambda(\bar{s},v)}} + \mathcal{O}(\delta x_i^2), \tag{8}$$

where $\overline{x_i}$ is the average entry frequency of alleles within a lineage. In the limit of zero effective selection coefficient ($\bar{s} \to 0$) with non-zero fluctuations ($v \neq 0$), the selection likelihood follows,

$$G_0(x|\overline{x_i};v) \equiv G(x|\overline{x_i};\bar{s}\to 0,v) = \frac{\overline{x_i}v(a_+ - a_-)}{\log\left|\frac{1-x/a_-}{1-x/a_+}\right|}, \tag{9}$$

which we use as a measure for neutral evolution with micro-evolutionary fluctuations. In the absence of fluctuations ($v \to 0$), the neutral propagator approaches $G_0(x|\overline{x_i}, v\to 0, \bar{s}\to 0) = x_i/x$, consistent with the evolutionary expectation for a single locus in neutrality [26].

The selection likelihood ratio follows,

$$g(x;\bar{s},v) = \frac{G(x|\overline{x_i};\bar{s},v)}{G_0(x|\overline{x_i};v)} = \frac{\bar{s}/v}{a_+ - a_-} \times \frac{\log\left|\frac{1-x/a_-}{1-x/a_+}\right|}{1-\left|\frac{1-x/a_+}{1-x/a_-}\right|^{\lambda(\bar{s},v)}} \tag{10}$$

.

The slope of the ratio $g(x;\bar{s},v)$ at small frequencies is $\bar{s}/2$, which provides an estimate for the effective selection strength. The micro-evolutionary fluctuations with an amplitude $v$ result in the flattening of the likelihood ratio at intermediate frequencies, as shown in Fig. 3 and Fig. S6. We infer the "effective selection pressure $\bar{s}$" and "selection fluctuations $v$" by fitting of the likelihood ratio function (eq. 10) to the estimates from lineages of different gene classes. Fig. S7 shows selection estimates for different BCR regions of lineages with distinct V-gene classes in all patients.

The time-ordered selection likelihood (interference likelihood) can be estimated using conditional probabilities under selection,

$$\begin{aligned} H(x|\overline{x_i};\overline{s_0},\overline{s_1},v) &= G(x|\overline{x_i};\overline{s_0},v)\,G(1|1-x;\overline{s_1},v) \\ &= \frac{\overline{s_0}\times\overline{x_i}}{1-\left|\frac{1-x/a_+}{1-x/a_-}\right|^{\lambda(\overline{s_0},v)}} \times \frac{1-\left|\frac{1-(1-x)/a_-}{1-(1-x)/a_+}\right|^{\lambda(\overline{s_1},v)}}{1-\left|\frac{1-1/a_-}{1-1/a_+}\right|^{\lambda(\overline{s_1},v)}}, \end{aligned} \tag{11}$$

where $\overline{s_0}$ and $\overline{s_1}$ are the effective selection strengths of the focal allele and the competing allele, respectively. In the absence of macro-evolutionary fluctuations, these effective selection strengths sum up to zeros, $\overline{s_0}+\overline{s_1}=0$.

The interference likelihood in neutrality ($\overline{s_0} \to 0$, $\overline{s_1} \to 0$) follows,

$$\begin{aligned} H_0(x|\overline{x_i}; v) &= G_0(x|\overline{x_i}; v)\, G_0(1|1-x, v) \\ &= \frac{v(a_+ - a_-)\,\overline{x_i}}{\log\left|\frac{1-x/a_-}{1-x/a_+}\right|} \times \frac{\log\left|\frac{1-(1-x)/a_-}{1-(1-x)/a_+}\right|}{\log\left|\frac{1-1/a_-}{1-1/a_+}\right|}. \end{aligned} \tag{12}$$

The interference likelihood ratio follows,

$$\begin{aligned} h(x; \overline{s_0}, \overline{s_1}, v) &= \frac{H(x|\overline{x_i}; \overline{s_0}, \overline{s_1}, v)}{H_0(x|\overline{x_i}; v)} \\ &= \frac{\overline{s_0}/v}{a_+ - a_-} \times \frac{\log\left|\frac{1-x/a_-}{1-x/a_+}\right|}{1 - \left|\frac{1-x/a_+}{1-x/a_-}\right|^{\lambda(\overline{s_0},v)}} \times \frac{1 - \left|\frac{1-(1-x)/a_-}{1-(1-x)/a_+}\right|^{\lambda(\overline{s_1},v)}}{\log\left|\frac{1-(1-x)/a_-}{1-(1-x)/a_+}\right|} \times \frac{\log\left|\frac{1-1/a_-}{1-1/a_+}\right|}{1 - \left|\frac{1-1/a_-}{1-1/a_+}\right|^{\lambda(\overline{s_1},v)}}. \end{aligned} \tag{13}$$

In the absence of long-term macro-evolutionary fluctuations ($\overline{s} \equiv \overline{s_0} = -\overline{s_1}$), the interference likelihood ratio $h(x)$ decays monotonically with allele frequency $x$. In this regime, if the focal allele is beneficial ($\overline{s} > 0$), the likelihood for a deleterious competing allele to take over and replace the focal allele becomes smaller as the frequency of the established focal allele $x$ increases within the lineage. In the case of a deleterious focal allele $\overline{s} < 0$, the likelihood for the focal allele to reach frequency $x$ in the first place becomes smaller as the frequency rises. Although short-term micro-evolutionary selection fluctuations could lead to a spontaneous beneficial state $-\overline{s} + \eta(t) > 0$ for otherwise a deleterious competing allele, these fluctuations are uncorrelated in time, and therefore, cannot maintain a positive selection long enough for the competing allele to establish within a lineage. Therefore, micro-evolutionary fitness fluctuations cannot boost the time-ordered likelihood under selection $h(x; \overline{s}, -\overline{s}, v)$ to exceed the neutral expectation.

Macro-evolutionary fluctuations for evolution with clonal interference lead to long-term turnover of the selection preference, and can convert a previously deleterious allele into a beneficial state. The full form of the interference propagator ratio (eq. 13) captures such effects by allowing two distinct effective selection strengths ($\overline{s_0}$, $\overline{s_1}$) for the focal and the competing alleles.

To characterize the fitness parameters, we fist infer the selection pressure on the focal allele $\overline{s_0}$ together with the amplitude of the micro-evolutionary fluctuations $v$ by fitting the selection propagator ratio $g(x; \overline{s_0}, v)$ (eq. 10) to the data. Using these estimates, we infer the effective selection on the competing allele $\overline{s_1}$ by fitting the interference propagator ratio $h(x; \overline{s_0}, \overline{s_1}, v)$ (eq. 13) to the data. It should be noted that $\overline{s_1}$ is the minimum strength of selection required for the competing allele to replace the focal allele and any stronger competing allele would result in a similar likelihood for interference. Fig. S6 shows such fits to different regions of BCRs for lineages from two distinct V-gene classes of patient 5. Fig. S9 shows deviations in the inferred selection strength of the competing alleles from the expected values in the absence of macro-evolutionary fluctuations.

**Fraction of selected sites.** Following the well established tradition of population genetics, we assume that synonymous mutations that do not change the amino acid provide a neutral gauge for evolution. In the case of frequency $x = 1$ the likelihood ratio $g(x)$ becomes equal to the ratio of the fixation probability $(D_n/P_n)\big/(D_s/P_s)$ where $D_n$ and $D_s$ are respectively the number of fixed non-synonymous and synonymous polymorphisms (i.e., substitutions) and $P_n$ and $P_s$ are total number of polymorphisms is each class. In other words, $g(x = 1)$ is equivalent to the McDonald-Kreitman test for selection based on the observed polymorphisms [22].

A selection likelihood ratio $g(x) = G(x)/G_0(x)$ larger than 1 implies an over-representation of non-synonymous compared to synonymous changes that reach frequency $x$ and is indicative of beneficial amino acid changes in a given region. For a total of $N$ non-synonymous mutations, we expect $NG_0(x)$ of these mutations to reach frequency $x$ by neutral evolution. The deviation between the observed number of non-synonymous mutations that reach frequency $x$, denoted by $n(x)$, and the neutral expectation quantifies the minimum fraction of beneficial mutations, $\alpha_{\text{benef.}}(x) = (n(x) - NG_0(x))/n(x) = (g(x) - 1)/g(x)$ [29]. On the other hand, a selection likelihood ratio smaller than 1 indicates negatively selected amino acid changes in a given region. The deviation from the expected number of non-synonymous mutations in neutrality, $NG_0(x) - n(x)$, is an estimate for the number of mutations that were suppressed due to deleterious fitness effects, indicating that at least a fraction $\alpha_{\text{del.}}(x) = 1 - g(x)$ of non-synonymous mutations to be under negative selection [29]. Similarly, we can compute the fraction of beneficial and deleterious mutations that are impacted by clonal interference, $\kappa_{\text{benef.}}(x) = (h(x) - 1)/h(x)$ for $h(x) > 1$, and $\kappa_{\text{del.}}(x) = 1 - h(x)$ for $h(x) < 1$.

**Inference of likelihood ratio statistics from data.** The descendants of a given mutation $\alpha$ on a lineage tree define the clade $\mathcal{C}^\alpha$. We calculate the frequency of mutation $\alpha$ appearing on a tree at time $t$, $x^\alpha(t)$, as the size of its clade $\mathcal{C}^\alpha(t)$ (i.e., the number of its observed descendent nodes) divided by the size of the lineage tree at time $t$. The time-trace for the frequency of a given mutation $x^\alpha(t)$ defines its evolutionary trajectory. The fraction of non-synonymous and synonymous mutations (e.g. within a certain gene group) that reach frequency $x$ during their history define the selection propagators $G(x)$ and $G_0(x)$, respectively. To infer statistically significant evidence for selection, we estimate propagators based on the mutations pooled from lineages of common gene classes, e.g. lineages with common V gene (Fig. 3b) or common V & J genes (Fig. 3c and Fig. S8). Table S1 reports the statistics of VJ-gene classes shown in Fig. 3c, Fig. S10.

It should be noted that the same mutation can occur independently on multiple backgrounds in a lineage and correspond to different strengths of selection in each case, depending on the time and the background that it has arisen on. Quantifying mutation frequencies based on their occurrences along the tree rather than from the alignment of the observed sequences within a lineage allows for adequate treatment of such independent events. Additionally, the maximum likelihood tree reconstruction algorithms could introduce spurious back mutations along the phylogeny. In our reconstructed lineages, however, the back mutations on average account for only a small fraction ($\sim 3\%$) of the mutations in a given lineage, and hence, have negligible impact on our analysis.

**Statistical analysis and error estimates of likelihood ratios.** We evaluate the expected error of a propagator at frequency $x$, by assuming binomial sampling from the total of $N$ non-synonymous and $N_0$ synonymous mutations (i.e., all the mutations observed in a given gene class). This results in the sampling errors $\sigma^2(x) = G(x)(1 - G(x))/N$ for non-synonymous and $\sigma_0^2(x) = G_0(x)(1 - G_0(x))/N_0$ for synonymous mutations, and a corresponding propagated error for the ratio, $g(x)$ in eq. 2. We use a similar approach to estimate the error for the interference likelihood ratio, $h(x)$ in eq. 3.

We use p-values based on the two-sample Kolmogorov-Smirnov test to compare selection statistics in different individual groups (i.e., untreated HIV patients, HIV patients with interrupted ART and healthy individuals) and in different BCR regions. Selection statistics among different regions of productive lineages are significantly distinct within each individual group (Fig. 3c, Fig. S10), with p-values $\ll 10^{-3}$ based on the two-sample KS test. In contrast, we do not infer a significant difference between selection patterns in CDR3 and FWR of unproductive lineages in healthy individuals (p-value $= 0.21$; Fig. S10). We use the pooled mutations statistics from both regions of unproductive lineages as the null expectation (dotted grey line in Fig. 3, Fig. S10). Selection statistics in productive lineages are distinct from the unproductive null with p-values estimated respectively in (CDR3, CDR1/2 and FWR) regions in untreated HIV patients ($4 \times 10^{-6}$, $0.$, $5 \times 10^{-10}$), in HIV patients with interrupted ART ($0.$, $0.$, $2 \times 10^{-5}$), and in healthy individuals (0, NA, $5 \times 10^{-3}$)[2]; see Fig. 3c, Fig. S4. Comparison between the two groups of HIV patients shows that their CDR3 statistics are significantly distinct (Fig. 3c) with p-value $= 5 \times 10^{-8}$ for selection statistics (Fig. 3c top) and p-value $= 5 \times 10^{-4}$ for clonal interference statistics (Fig. 3c bottom). Such a shift is not present for mutations in CDR1/2 and FWR (p-values $> 0.1$).

[2] p-values =0. are estimates within the machine error.

#### 5 Robustness of selection statistics

**Heterogenous BCR hypermutation rates.** It should be noted that the heterogenous and context dependent somatic hyper-mutation rates during affinity maturation [30–34] introduce BCR-specific biases that could influence inference of selection. In order to verify the robustness of our method, we have simulated the process of affinity maturation based on two distinct BCR-specific hyper-mutation models along the inferred BCR lineage phylogenies: (i) S5F hypermutation profile [30] and (ii) IGoR profile [34]; see the paragraph on Simulations for details.

First, we characterize the effect of hypermutation biases on the signal of selection based on the SFS. In simulated trees, we observe a similar shape for the SFS polarized on mutations in different regions of BCRs (Fig. S4), in contrast to the region-specific signal for selection observed from the real data in Fig. 2. The upturn in the high frequency tail of the SFS is a property of mutation propagation along a phylogeny and is closely related to the skewness and existence of long branches in a phylogeny. In our simulations, we introduce mutations from context-dependent hypermutation models along the BCR phylogenies inferred from real data (see the paragraph on simulations). Therefore, the high-frequency upturn of the SFS is sustained but not the identity of the mutations associated with this signal (Fig. S4). Moreover, the shape of the shuffled SFS does not depend on the details of the hyper-mutation model.

In addition, Fig. S10a, b show that the region-specific likelihood ratios $g(x)$, $h(x)$ are insensitive to the heterogenous hypermutation statistics and such biases do not produce spurious evidence for selection and clonal interference. The biases in the BCR profiles introduce hotspots in CDR regions that are enriched in volatile codons, and hence, can have high rates of amino acid mutations [35, 36]. Our simulations of neutral affinity maturation captures the impact of such hotspots. The selection likelihood statistics in Fig. 3 show a significantly larger over-representation of non-synonymous to synonymous mutations in CDR regions of real BCR data compared to the simulated trees (Fig. S10), indicating that mutational hotspots alone cannot explain the sequence statistics observed in CDRs. Moreover, the likelihood ratio is insensitive to the initial frequency of an allele within a lineage [23] and provides a robust measure for inference of selection in evolving genealogies.

**Sequencing depth of BCR repertoires.** The data from BCRs of healthy individuals from ref. [4] has a higher depth than the data from HIV patients [1]. To evaluate the robustness of our statistics, we repeat our analysis on the heathy BCR repertoire data under-sampled to the level of the sequencing depth in HIV patients. Fig. S2 shows that the distribution for the terminal branch length statistics is insensitive to the sequencing depth. In Fig. S8 we compare the selection likelihood statistics inferred from the full data to the under-sampled data in healthy individuals. In both cases we see an over-representation of positively selected amino acid changes in the CDR3 compared to the FWR of BCRs. In the under-sampled data, we infer a significantly weaker strength of positive selection on amino acid changes in CDR3, with an averaged fraction of $\overline{\alpha}_{\text{benef.}} = 22\%$ for beneficial and $\overline{\alpha}_{\text{del.}} = 6\%$ for deleterious changes, in comparison to the estimates from the full dataset in Table 1; KS-test p-value between distributions of the sub-sampled and the full dataset for the fraction of selected amino acid changes are, $5.0 \times 10^{-6}$ for beneficial and $0.04$ for deleterious changes. In addition, we infer a significantly lower fraction of deleterious changes in FWR in the subsampled data compared to the full dataset, with an average $\overline{\alpha}_{\text{del.}} = 16\%$ and KS-test p-value= $4.9 \times 10^{-6}$, but a comparable fraction of beneficial changes $\overline{\alpha}_{\text{benef.}} \sim 7\%$. Overall, we infer that subsampling from the data results in less stark differences between the inferred selection strengths across different BCR regions, and hence, it results in a conservative estimate for the strength of positive selection in CDR3.

#### 6 Simulations

**Simulated trees.** In Fig. 2b, we compare the branch length characteristics of the BCR genealogies (in units of coalescence time) with the neutral expectation from 2000 simulated trees with Kingman's coalescent, generated by the beta coalescent algorithm with parameter $\alpha = 1$ [37]. The sizes of the simulated trees in the neutral ensemble are drawn from the BCR lineage size distribution in HIV patients. The schematic trees in Fig. S2a are also generated similarly by the beta coalescent algorithm [37] with parameters $\alpha = 1$ for neutral evolution and $\alpha = 2$ for rapid adaptation.

**Bootstrapping site frequency spectrum.** We characterize the uncertainty in the neutral site frequency spectrum from 100 realizations of simulated neutral trees (Kingman's coalescent), which we generate by the beta coalescent algorithm with parameter $\alpha = 1$ [37]. For each dataset (i.e., HIV patients, healthy-productive and healthy-unproductive), a simulated realization contains an equal number of trees with the same tree sizes as the actual data. The uncertainty in the SFS varies across datasets (see Fig. 2, Fig. S3) since the SFS is an aggregate measure across mutations in all trees of a given dataset and the datasets differ in the number of trees, and hence, the number of mutations (see Table S1). For a given SFS $f(\nu)$, we evaluate the test statistics $\delta f = f(\nu \to 1) \big/ \min[f(\nu)]$ to quantify the observed high-frequency upturn in the real data and also its uncertainty from the simulated neutral realization. We estimate the p-value for the observed $f(\nu)$ from the distribution of the test statistics across the corresponding simulated realization.

**Null model for context-dependent affinity maturation.** We simulate mutations along BCR lineage trees according to two context-dependent models of hyper-mutation, (i) IGoR statistics [34] and (ii) S5F [30]. For a given branch on a lineage tree, we draw a number of mutations equal to the branch length from a multinomial distribution with position-specific weights determined by the hypermutation models. Due to the changes in the sequence, we update the position weights at each internal node of the tree to account for context-dependent hyper-mutation rates. This procedure reshuffles the identity of mutations along BCRs according to the neutral hyper-mutation models, while preserving the shape of tree. Fig. S4 and Fig. S10 show that SFS and the propagator statistics do not recover evidence for region-specific selection in BCRs in the simulated lineages. Therefore, the original selection signal in Figs. 2, 3 are not reflecting any spurious effect due to heterogenous mutation rates.

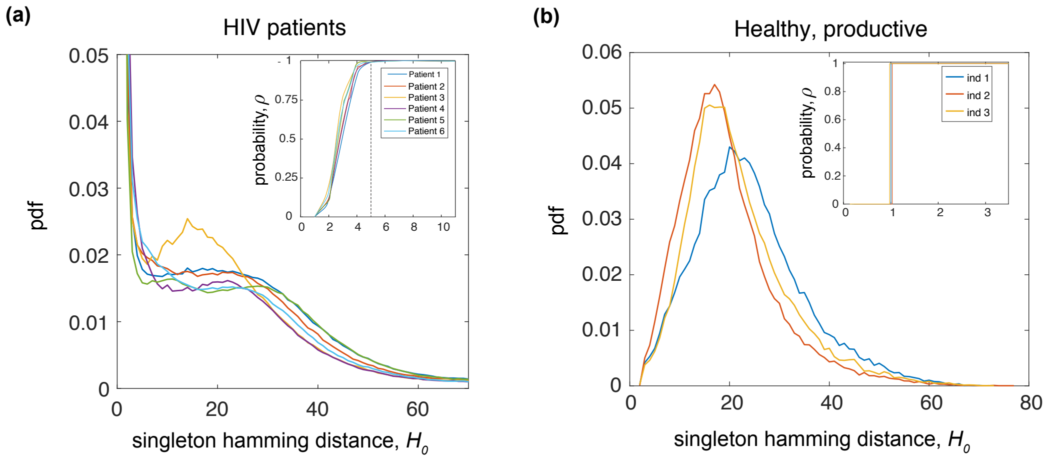


**Figure S1: Singletons in the BCR repertoire.** The probability density for the minimum Hamming distance of singleton (a read with multiplicity 1) to any non-singleton, $H_0$, is shown for **(a)** each HIV patient and **(b)** each healthy individual. We use a hidden Markov model and decompose the observed probability densities $W(H_0)$ into a mixture of a Poisson distribution, related to sequencing errors, and a Gamma distribution, as the true distribution of $H_0$ in each individual. The insert in each panel shows the sigmoid-form probability function $\rho$ for a singleton with a Hamming distance $H_0$ to be a true independent BCR sequence and not to be generated by sequencing errors. In our analysis, we only admit sequences with low uncertainty $\rho(H_0) \simeq 1$, corresponding to $H_0 \geq 5$ in HIV patients and $H_0 \geq 1$ in healthy individuals.

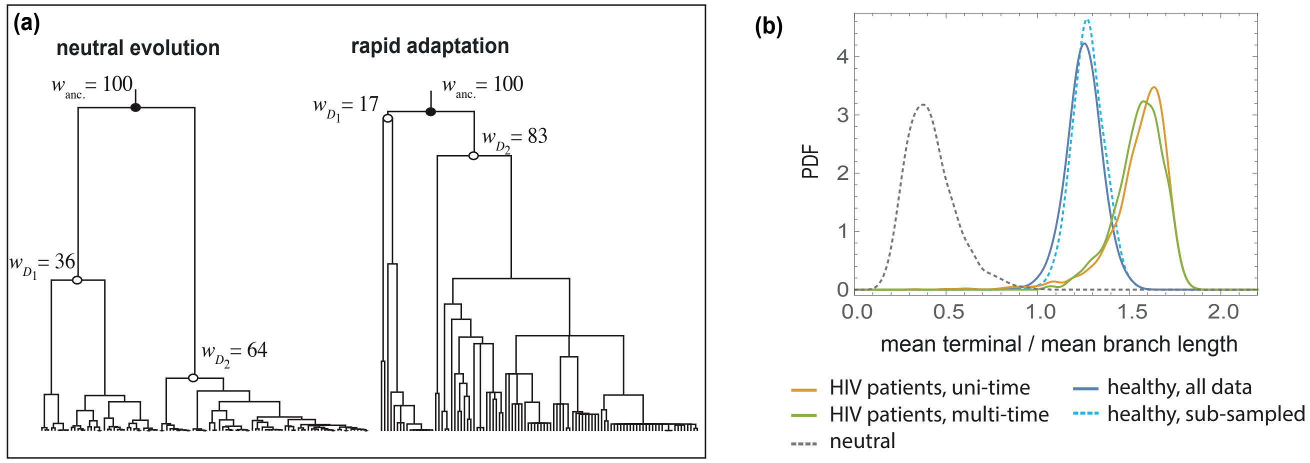


**Figure S2: Impact of selection on lineage tree statistics. (a)** Simulated phylogenies for neutral evolution (left) and rapid adaptation with positive selection (right), generated by the coalescence package [37] and plotted with FastTree [9]. The weights of the ancestral node $w_{\text{anc.}}$ and its daughters $w_{D_1}, w_{D_2}$ (i.e., their clone size) are indicated in each phylogeny. **(b)** shows the distribution of mean terminal branch length (in units of coalescence time) relative to the mean length of all branches in sub-sampled BCR lineages. The probability density of branch length statistics is comparable between BCR lineages of HIV patients that are present in only a single time point (orange) and lineages sampled over multiple time points (green). The shape of the distribution is insensitive to the sequencing depth of the BCR repertoires. We sub-sampled the BCR data in healthy individuals to the level of sequencing depth in HIV patients. The probability density for the branch length statistics of lineages reconstructed from the sub-sampled repertoires in healthy individuals (dotted light blue) is comparable to the statistics from the complete data (dark blue). The corresponding distribution for the simulated neutral trees (similar to Fig. 2b) is shown by a dotted gray line. The elongated terminal branches in BCR lineages is indicative of positive selection.

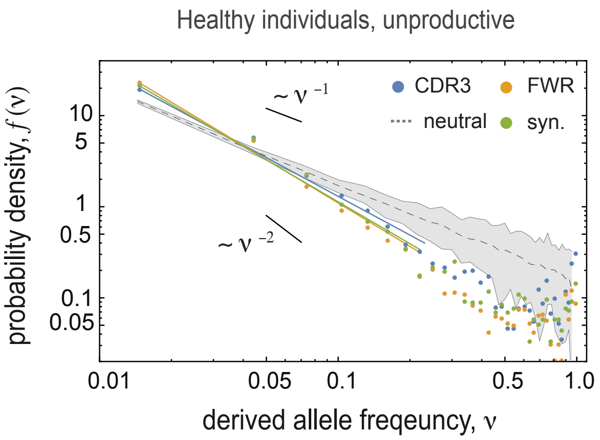


**Figure S3: Site frequency spectrum for unproductive lineages.** SFS $f(\nu)$ is shown for mutations in different regions of BCRs (distinct colors) in unproductive lineages of size $> 100$ pooled from healthy individuals (total of 131 lineages). The grey area shows the span of SFS across 100 realizations of 131 simulated neutral trees (Kingman's coalescent) with sizes equal to the unproductive BCR lineages; see the simulation section and Fig. 2c for SFS in productive lineages.

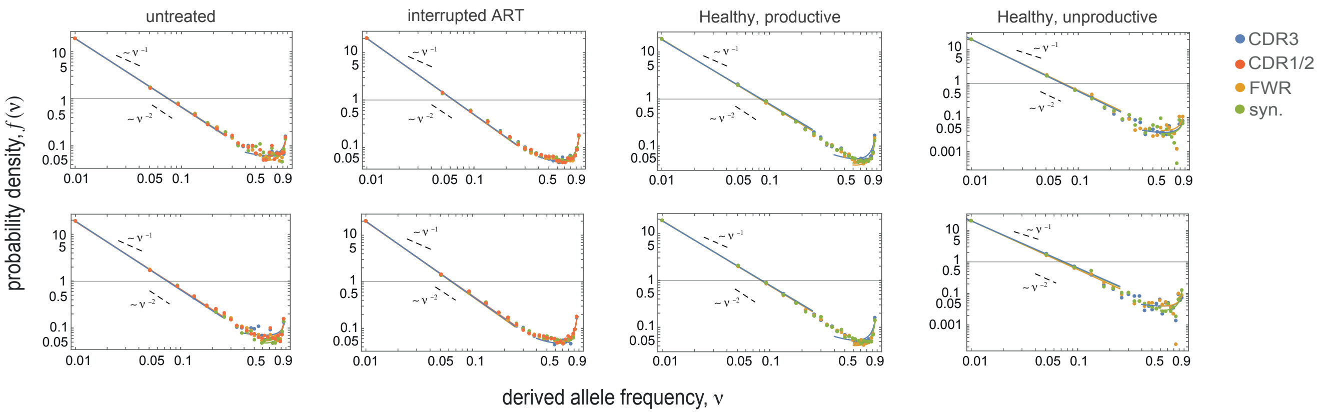


**Figure S4: Robustness of SFS to BCR hypermutation biases.** The figure shows the site frequency spectra for simulated neutral hypermutation processes along the inferred B-cell lineages, as described in the Methods. We use two hypermutation models, IGoR statistics [34] (top row) and the S5F model [30] (bottom row) for each panel. The SFS estimated for lineages with randomized mutations show homogenous patterns across different BCR regions in all categories of individuals and BCRs, indicating that the region-specific upturn of SFS in Fig. 2 is not an artifact caused by the BCR hypermutation biases. SFS is shown for mutations in different regions of BCRs (colors) in healthy individuals, HIV patients with interrupted ART and in untreated HIV patients.

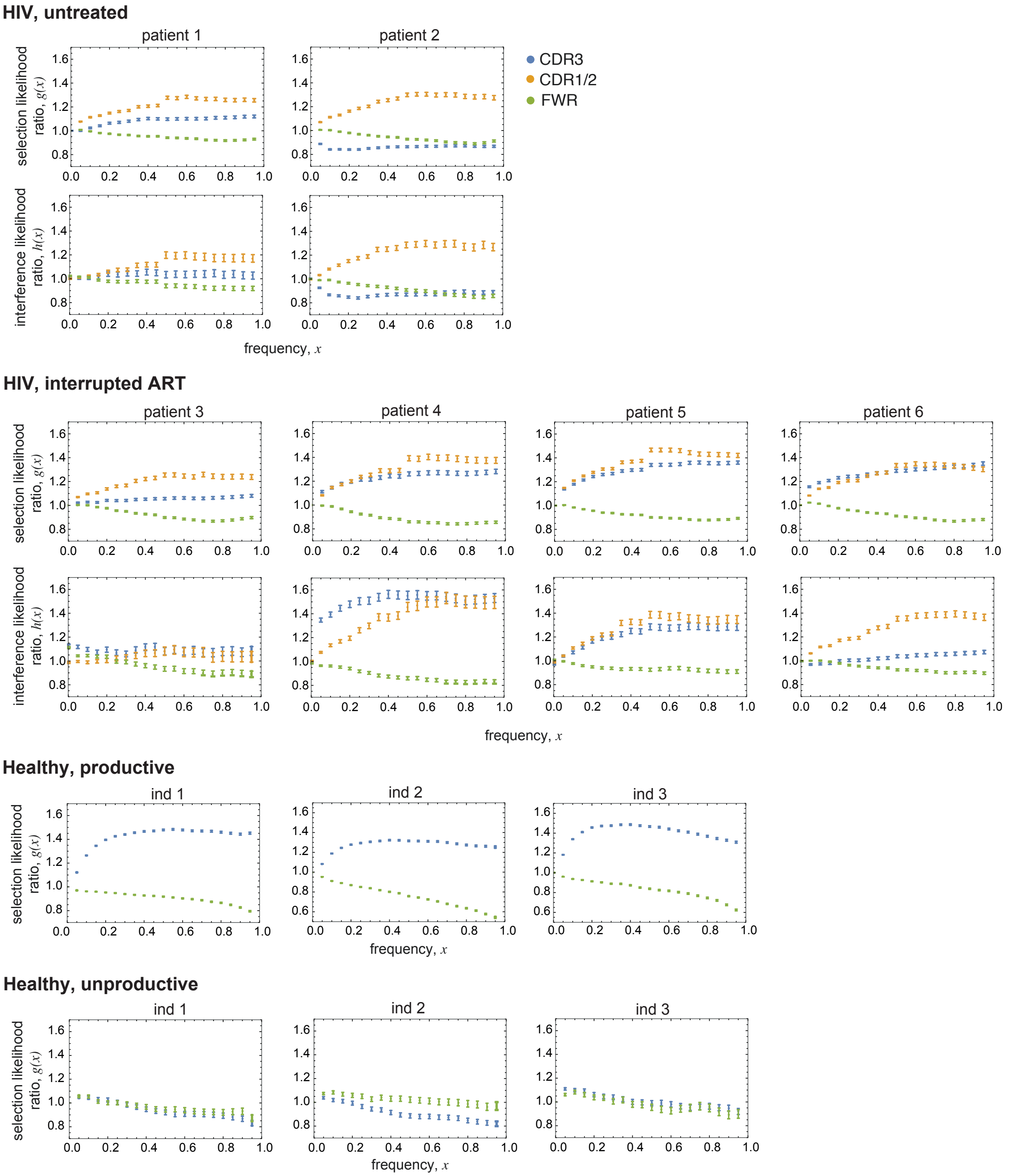


**Figure S5: Selection and interference likelihood ratios in all individuals.** Panels show selection and interference likelihood ratios $g(x)$, $h(x)$ in HIV patients (untreated and with interrupted ART) and the selection likelihood ratio $g(x)$ in productive and unproductive lineages of healthy individuals, aggregated over all lineages in each individual. We consistently see strong evidence for negative selection in FWR regions of productive lineages and positive selection in both or either of CDR regions. We do not see such distinction in unproductive lineages. Note that the repertoire level averaged likelihood ratios are highly coarse grained statistics and miss the gene-specific evidence for selection and clonal interference, as shown in Fig. 2.

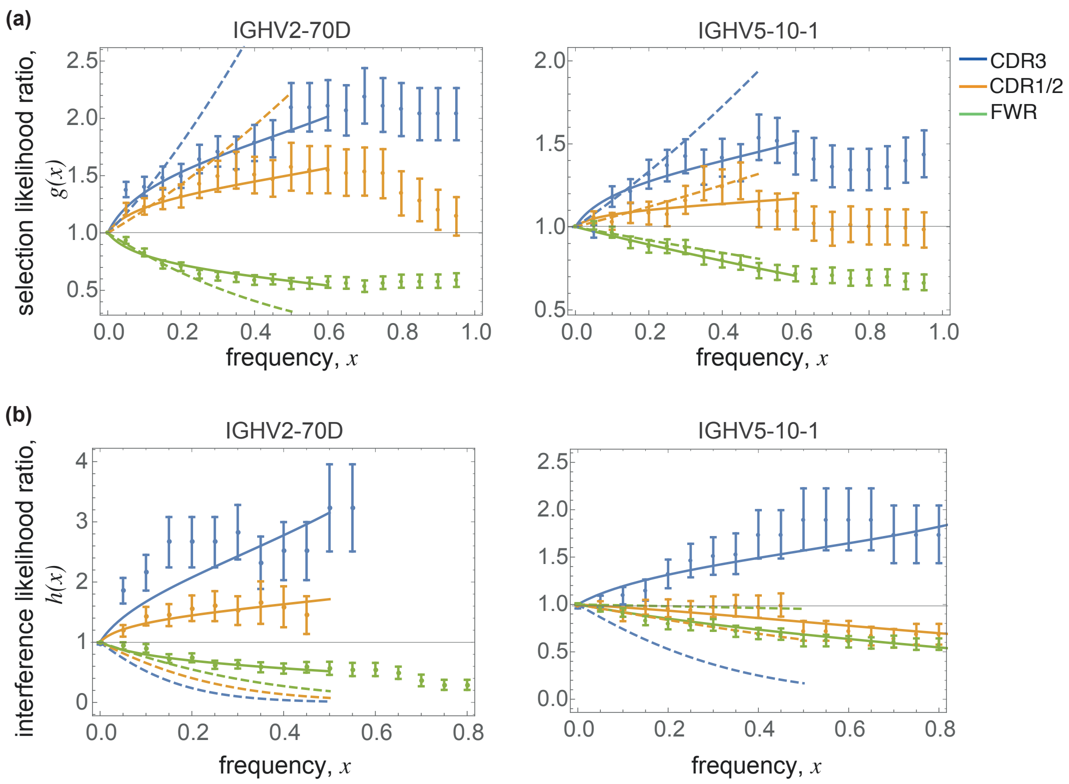


**Figure S6: Selection and clonal interference in BCR lineages.** **(a)** shows theoretical fits to selection likelihood ratio $g(x; \overline{s}, v)$, based on the selection model with micro-evolutionary fluctuations $v > 0$ (full lines) and without any fluctuations ($v = 0$) (dashed lines) in different BCR regions (colors) for lineages from Patient 5 with two distinct V-gene classes IGHV2-70D (left) and IGHV5-10-1 (right); see Methods and (eq. 10). In both cases we infer strong effective selection coefficients $\overline{s} = [14.0 \pm 2,\ 10.1 \pm 2.5,\ -7.2 \pm 0.5]$ (left) and $\overline{s} = [7.5 \pm 1.6,\ 3.9 \pm 2.2,\ -1.2 \pm 0.1]$ (right), in CD3, CDR1/2 and FWR respectively. The data is inconsistent with the evolution with constant selection (dashed lines), suggesting that micro-evolutionary fitness fluctuations are necessary to explain the flattening of the likelihood ratio, up to intermediate frequencies (full lines). **(b)** shows the theoretical fits to the interference likelihood ratio $h(x; \overline{s_0}, \overline{s_1}, v)$ (full lines), where the effective selection on the competing allele $\overline{s_1}$ is the fitted coefficient and the strength of selection on the focal allele $\overline{s_0} \equiv \overline{s}$ (in (a)) and the fluctuation amplitude $v$ are the estimates from (a); see Methods and eq. 13. The prevalence of macro-evolutionary fitness fluctuations is evident from the data presented in all panels: At high frequencies ($x > 0.6$), the extrapolation of the fits with micro-evolutionary fluctuations (full lines) deviate from the data in (a), as the role of macro-evolutionary fluctuations become crucial in describing the distribution of alleles within a lineage (Methods). In (b), the data is inconsistent with the expected interference likelihood ratio in the absence of macro-evolutionary fluctuations $\overline{s_1} = -\overline{s_0}$ (dashed line), suggesting an evolutionary regime with strong clonal interference for BCR maturation.

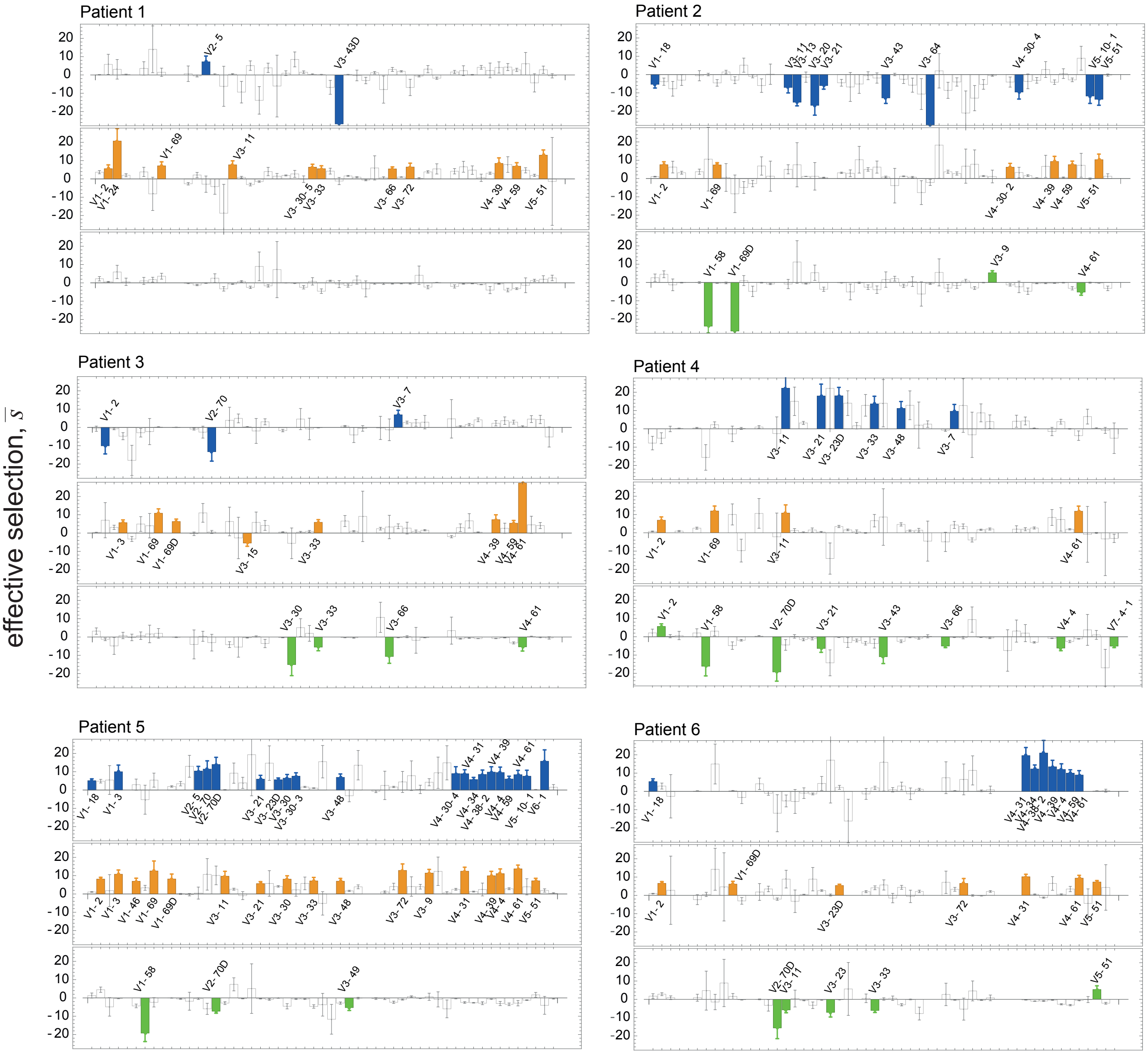


**Figure S7: Inferred selection strengths in different patients.** Effective selection strength $\overline{s} = 2N\overline{\sigma}$ (in units of $N$) is inferred by fitting the theoretical expectation in the presence of fluctuations (eq. 10) to the selection likelihood ratio estimated from lineages in different V-gene classes in each patient. In each patient panel, the inferred selection strengths are reported separately for the distinct BCR regions, CDR3 (blue, top box), CDR1/2 (orange, middle box) and FWR (green, bottom box). In each region, the V-genes for which the selection strengths are substantial $|\overline{s}| > 5$ and are confidently inferred (p-value $\leq 0.01$ of non-linear fit) are indicated in the chart, with error bars indicating the confidence interval for the inferred selection coefficients. The selection strength for other present V-gene classes are indicated with empty boxes and corresponding confidence intervals. V-genes are ordered alphabetically along the $x$-axis in all panels.

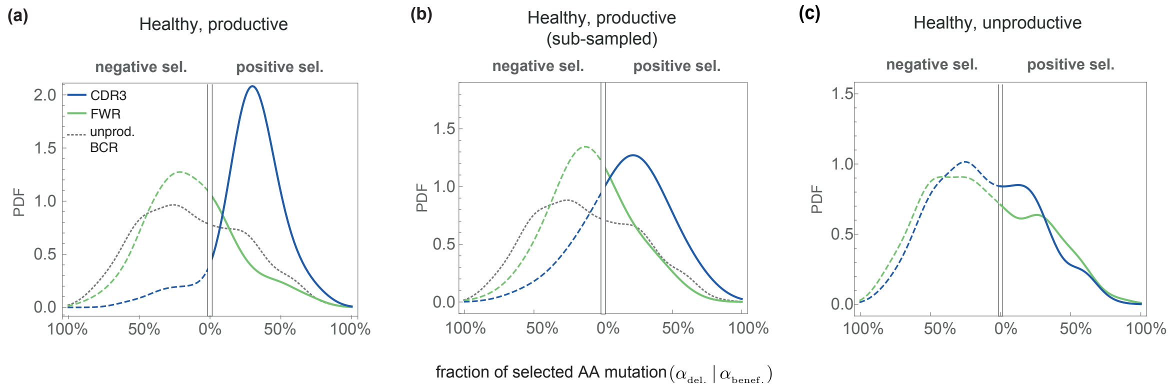


**Figure S8: Fraction of selected BCR mutations in healthy individuals.** The probability density across distinct VJ-gene classes for the (minimum) fraction of beneficial (right) $\alpha_{\mathrm{benef.}}$ and deleterious (left; inverted x-axis) $\alpha_{\mathrm{del.}}$ amino acid changes that reach frequency $x = 80\%$ within a lineage is shown for different regions of BCRs in healthy individuals for **(a)** all productive lineages **(b)** productive lineages in repertoires sub-sampled to the level of sequencing depth in HIV patients and **(c)** unproductive lineages. Similar to HIV patients (Fig. 3), the CDR3 mutations in productive lineages of healthy individuals are under positive selection, whereas the FWR mutations are under negative selection. We do not infer any significant difference between selection patterns in the CDR3 and in the FWR region of unproductive lineages (p-values $= 0.21$, two-sample KS-test). The probability density for mutations pooled from both regions of unproductive lineages is shown in **(A, B)** as the null expectation (dotted grey line), similar to Fig. 3.

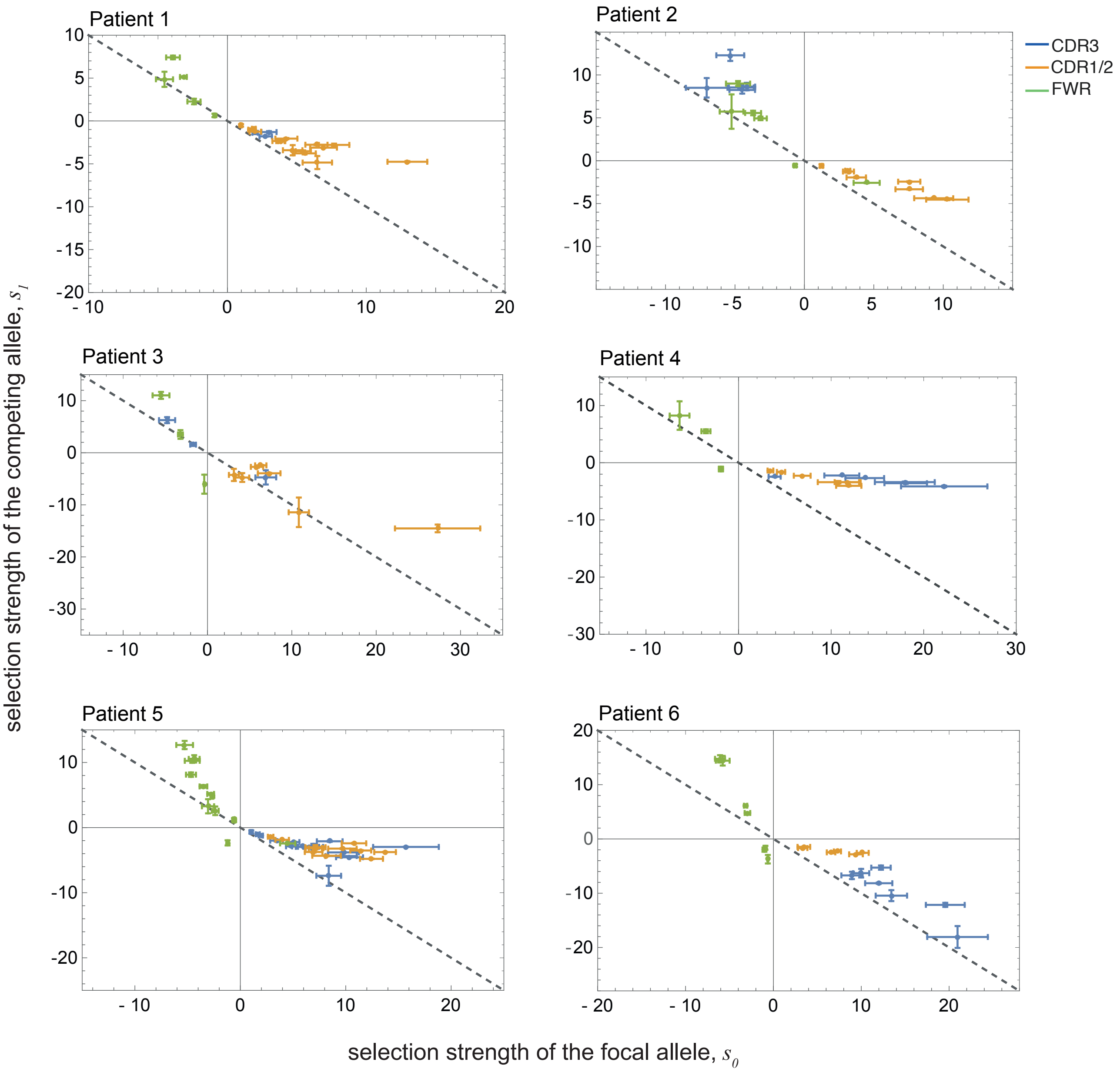


**Figure S9: Prevalence of macro-evolutionary fitness fluctuations in different patients.** Selection strengths for the focal allele $\overline{s_0}$, inferred from the selection likelihood ratio in V-gene classes $g(x; \overline{s_0}, v)$ (eq. 10), is compared to the strength of selection for the competing allele $\overline{s_1}$ inferred from the interference likelihood ration $h(x; \overline{s_0}, \overline{s_1}, v)$ (eq. 13) in all patients (panels). In each patient panel, the confidently inferred selection strengths (p-value $\leq 0.05$ of the non-linear fit to the likelihood functions are shown for distinct BCR regions (colors). The diagonal dashed line in each panel indicates the expectation for evolution without macro-evolutionary fluctuations, with $\overline{s_1} == \overline{s_0}$. The higher strength of selection in the competing alleles $\overline{s_1} > -\overline{s_0}$, observed clearly in patients 4-5, signal the persistence of macro-evolutionary fluctuations in fitness, and hence, a strong clonal interference within BCR lineages, consistent with the results in Fig. 4b. Error bars indicate the confidence interval for the inferred selection coefficients.

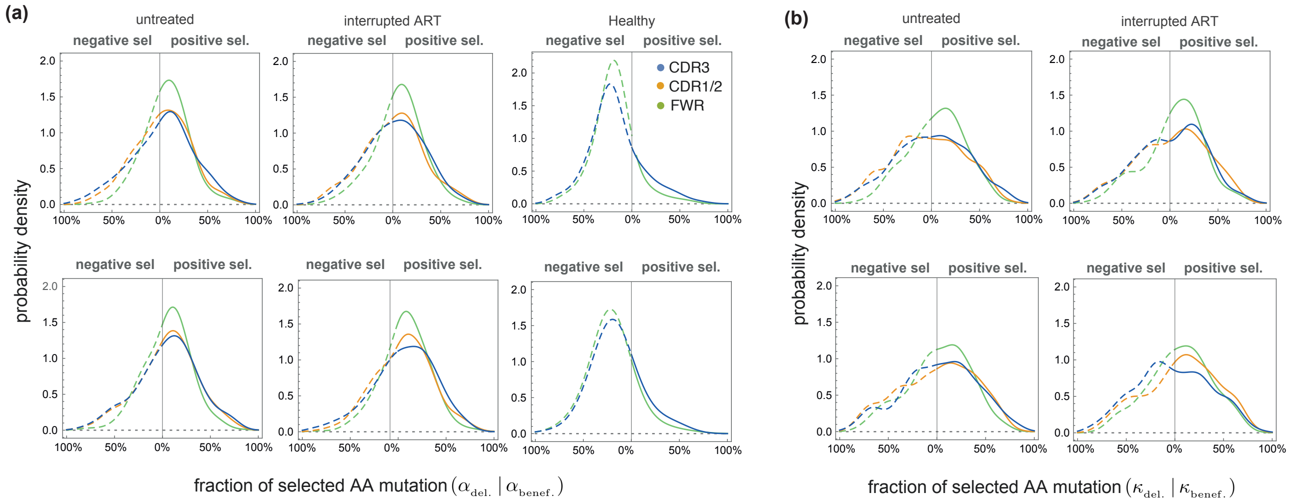


**Figure S10: Robustness of selection inference to BCR hypermutation biases.** (**a**) Selection and **(b)** interference likelihood ratios are shown for lineages with randomized mutations, using two hypermutation models (Methods), IGoR statistics [34] (top row) and the S5F model [30] (bottom row). Similar to Fig. 3, each panel shows the probability density across distinct VJ-gene classes for **(a)** the minimum fraction of beneficial $\alpha_{\mathrm{benef.}}$ and deleterious $\alpha_{\mathrm{del.}}$ mutations that reach frequency $x = 80\%$, and (**b**) for beneficial $\kappa_{\mathrm{benef.}}$ and deleterious $\kappa_{\mathrm{del.}}$ mutation fractions that reach frequency $x = 60\%$ and later go extinct. These statistics are estimated for mutations in different regions of BCRs (colors) in healthy individuals, HIV patients with interrupted ART and in untreated HIV patients. In contrast to the real data (Fig. 3), we do not observe any region-specific pattern of selection for simulated lineages with context-dependent hypermutation models, indicating that the inferred region-specific pattern of selection (Fig. 3) is not an artifact of biased hypermutation statistics in BCRs.

| | | HIV infected (untreated ) | | HIV infected (interrupted ART at week 48) | | | | Healthy | | |
|---|---|---|---|---|---|---|---|---|---|---|
| | | patient 1 | patient 2 | patient 3 | patient 4 | patient 5 | patient 6 | ind 1 | ind 2 | ind 3 |
| **productive BCRs** | | | | | | | | | | |
| # unique BCR sequences | | 148,404 | 159, 920 | 118,417 | 103,623 | 273,766 | 226,530 | 998,386 | 803,974 | 723,746 |
| # of lineages | size | | | | | | | | | |
| | $> 20$ | 3,343 | 3,700 | 3,154 | 2,112 | 4,209 | 4,126 | 21,492 | 15,726 | 15,781 |
| | $> 50$ | 785 | 773 | 613 | 485 | 1,246 | 1,205 | 5,242 | 3,978 | 3,390 |
| | $> 100$ | 207 | 198 | 101 | 125 | 482 | 411 | 1,525 | 1,270 | 1,001 |
| # of VJ-gene classes | | 192 | 181 | 144 | 195 | 176 | 146 | 260 | 231 | 247 |
| **unproductive BCRs** | | | | | | | | | | |
| # unique BCR sequences | | 0 | 0 | 0 | 0 | 0 | 0 | 33,982 | 43,930 | 35,276 |
| # of lineages | size | | | | | | | | | |
| | $> 20$ | 0 | 0 | 0 | 0 | 0 | 0 | 899 | 1041 | 963 |
| | $> 50$ | 0 | 0 | 0 | 0 | 0 | 0 | 177 | 198 | 155 |
| | $> 100$ | 0 | 0 | 0 | 0 | 0 | 0 | 29 | 57 | 45 |
| # of VJ-gene classes | | 0 | 0 | 0 | 0 | 0 | 0 | 181 | 47 | 189 |
| time samples (weeks) | | 0, 4, 16, 24, 52, 72, 120 | 0, 4, 12, 16, 24, 52 60, 108 | 4, 12, 16, 24, 52, 60, 108 | 4, 12, 16, 24, 52, 60, 108 | 4, 12, 16, 24, 52, 60, 108 | 0, 4, 16, 24, 52, 60, 108 | 0 | 0 | 0 |

**Table S1: Statistics of BCR repertoires.** The table presents the number of uniques sequences used to reconstruct lineages with size ($> 20$), the number of reconstructed lineages with size ($> 20$), ($> 50$) and ($> 100$), the number of inferred VJ-gene classes, and the sampled time points after the start of the study in HIV infected patients and in healthy individuals. BCR sequences extracted from healthy individuals do not cover the CDR1/2 region.

| wks since seroconversion / count (ml$^{-1}$) | | wk 0 | wk 4 | wk 12 | wk 16 | wk 24 | wk 52 | wk 60 | wk 108 |
|---|---|---|---|---|---|---|---|---|---|
| Patient 1 | viral load | $>$ 500 k | 279876 | 376588 | NA | $>$ 500k | 160704 | 198046 | 110382 |
| | CD4+ | 530 | 470 | 640 | NA | 440 | 480 | NA | 470 |
| Patient 2 | viral load | $>$ 500 k | 271878 | NA | 138602 | 117512 | 89926 | 88003 | 147536 |
| | CD4+ | NA | 550 | 600 | 690 | 630 | 530 | 440 | NA |
| Patient 3 | viral load | 129037 | 709 | $<$ 50 | 52 | $<$ 50 | 489 | 6005 | 20459 |
| | CD4+ | 870 | 1610 | 1130 | 1230 | 1180 | 1200 | 870 | 1460 |
| Patient 4 | viral load | 99046 | 206 | $<$ 50 | $<$ 50 | $<$ 50 | 349343 | 7922 | 39868 |
| | CD4+ | 550 | 830 | 1130 | 620 | 850 | 500 | 650 | 510 |
| Patient 5 | viral load | 67580 | 722 | 122 | $<$ 50 | $<$ 50 | 269443 | 4247 | 196326 |
| | CD4+ | 520 | 490 | 620 | 550 | 480 | 600 | 740 | 500 |
| Patient 6 | viral load | 315764 | 620 | $<$ 50 | 187 | NA | 101221 | 339135 | 76860 |
| | CD4+ | 550 | 770 | 900 | 390 | NA | 660 | 320 | NA |

**Table S2: Viral load and CD4+ counts in HIV patients.** Data collected from HIV patients since seroconversion at week 0. Patients 1 and 2 were untreated, while patients $4-6$ received ART until week 48.
